# Enhancing Gene Expression Representation and Drug Response Prediction with Data Augmentation and Gene Emphasis

**DOI:** 10.1101/2024.05.15.592959

**Authors:** Diyuan Lu, Daksh P.S. Pamar, Alex J. Ohnmacht, Ginte Kutkaite, Michael P. Menden

## Abstract

Representation learning for tumor gene expression (GEx) data with deep neural networks is limited by the large gene feature space and the scarcity of available clinical and preclinical data. The translation of the learned representation between these data sources is further hindered by inherent molecular differences. To address these challenges, we propose GExMix (**G**ene **Ex**pression **Mix**up), a data augmentation method, which extends the Mixup concept to generate training samples accounting for the imbalance in both data classes and data sources. We leverage the GExMix-augmented training set in encoder-decoder models to learn a GEx latent representation. Subsequently, we combine the learned representation with drug chemical features in a dual-objective enhanced gene-centric drug response prediction, i.e., reconstruction of GEx latent embeddings and drug response classification. This dual-objective design strategically prioritizes gene-centric information to enhance the final drug response prediction. We demonstrate that augmenting training samples improves the GEx representation, benefiting the gene-centric drug response prediction model. Our findings underscore the effectiveness of our proposed GExMix in enriching GEx data for deep neural networks. Moreover, our proposed gene-centricity further improves drug response prediction when translating preclinical to clinical datasets. This highlights the untapped potential of the proposed framework for GEx data analysis, paving the way toward precision medicine.

## 1 Introduction

Deep learning capabilities have been explored in many fields with impressive results. Numerous studies have shown that encoder-decoder-based models can learn meaningful latent representations from GEx data [1, 2, 3]. However, leveraging the learned representations for clinically relevant tasks remains challenging. In human cancer patients, who exhibit diverse responses to treatments given the individual molecular tumor characteristics, personalized treatment approaches are necessary for improved outcomes [4]. Thus, developing drug sensitivity predictors for cancer patients encounters challenges, including limited patient-drug pair data, patient profile heterogeneity, and unknown biochemical factors.

Recent advancements in obtaining molecular features, such as gene expression, at a faster speed and lower costs have motivated the use of comprehensive tumor molecular profiles to tailor cancer treatments to individual tumors [5]. The Cancer Genome Atlas (TCGA) has significantly contributed to extensive clinical human genomic data repositories across a wide variety of cancer types [6].

These advancements have been further accelerated with the increasing availability of tumor data in preclinical data sources, e.g., animal studies, and cell lines [7], such as Genomics of Drug Sensitivity in Cancer (GDSC) [8] and Cancer Cell Line Encyclopedia (CCLE) [9].

Despite these advancements, transferring drug response insights from cell lines to human patients remains challenging. The reasons include the limited amount of available human tumor data, limited representativeness of minority data classes (cancer types) with a small number of available cell lines, and inherent molecular profile differences between cell lines and tumors due to the biological and technical confounders [10]. These factors hinder the translation of predictions from preclinical studies like the GDSC and CCLE to clinical settings represented by the TCGA [11].

There have been studies aiming to incorporate knowledge from both preclinical and clinical GEx datasets for improving the potential knowledge transfer from cell lines to human patients. These approaches include explicit source information disentanglement training [10], domain-generalization [3] treating human tumor dataset as out-of-domain targets, or few-shot learning with samples from target domains [12]. However, these approaches are constrained by the limited number of GEx samples and do not explore the potential of data augmentation for enriching the training set, thus, failing to utilize the available data. When translating drug response to the clinical context, the drug response prediction (DRP) is often evaluated in a one-model-per-drug fashion in a small subset of available drugs, thus lacking generalized insights for unseen drugs.

To address these challenges, we propose GExMix, a data augmentation method to improve general representation learning with GEx data. In this study, we focus on GEx data from two sources, i.e., human tumors and cell lines. While data augmentation is common practice in various tasks, it remains underexplored for GEx data. Inspired by Mixup [13], GExMix extends the concept to account for the imbalance in cancer types and data sources, i.e., human tumors and cell lines. The rationale is that it can generate more samples from minority cancer types as well as minority data sources. The GExMix augmented training set is subsequently employed to train the encoder-decoder framework, aiming to learn the latent representation of GEx data from diverse sources. Furthermore, we propose a dual-objective approach for the drug response predictor to emphasize the contribution of GEx features in DRP while leveraging the above-learned representation. The objective consists of a classification loss for DRP and a mean-square-loss to reconstruct the input GEx embeddings with the output from the same layer. This design strategically preserves the GEx information for DRP and the predictor is compelled to excel in both DRP and upholding fidelity of the input gene expression representation.

Our contributions can be summarized as follows. 1. We propose GExMix, a data augmentation method, for enhancing the representation learning with GEx data. 2. We propose a gene-centric dual-objective DPR training framework to emphasize the contribution of gene expression in a unified model for multi-drug response prediction. 3. We rigorously validate the effect of GExMix and gene-centric DRP learning across several deep encoder-decoder-based models and ablation experiments. 4. We demonstrate improvement in overall latent embeddings and generalization performance with our proposed methods, providing a compelling proof-of-concept and showcasing promising transferability to clinical patient DRP.

## 2 Related Work

### Autoencoders with gene expression

Gene expression (GEx) levels, reflecting the transcription of DNA into messenger RNA (mRNA) for protein synthesis, are affected by factors including epigenetic factors, gene mutations, tissue origins, and disease conditions [14]. Constrained by the vast number of features and a limited number of samples, many studies have proposed to learn an abstract representation of GEx data [1, 2, 10]. Autoencoders exhibit great potential in capturing complex non-linear correlations in the input data in an unsupervised manner [1, 2]. They allow the investigation of GEx data at a more abstract level. For example, in [15], the authors proposed a deconfounding autoencoder for learning robust gene expression embeddings, which is achieved by enforcing an adversarial loss on data sources. The context-aware deconfounding autoencoder (CODE-AE) [10], designed to learn an aligned representation of GEx data with shared and private encoders for two sources, i.e., TCGA and CCLE, aiming to transfer source-dependent knowledge from cell lines to clinical drug responses. However, methods to improve the latent representation of GEx data are underexplored in these studies. We adopt the encoder design from [10], while focusing on investigating the impact of our proposed data augmentation, GExMix, on GEx representation learning for subsequent precision medicine applications.

### Mixup augmentation

Mixup [13] was proposed for augmenting training samples to improve the generalization ability of DNNs and has been widely used in other fields, such as audio [16] and text [17]. There have been variants like CutMix [18], mixing with cut augmentation, and Manifold mix [19], mixing the hidden states from randomly selected layer depths. Investigations on efficacy, robustness, and generalization [20], as well as its impact on calibration for DNNs have been explored [21, 22]. Our proposed GExMix adapts Mixup and tailors it for GEx data analysis in precision oncology, specifically addressing source imbalance and potential biological and technical confounders.

### Drug response prediction

The development of DRP models is a central pillar for enabling personalized treatments [23]. In recent years, numerous methods have been proposed and benchmarked, showcasing their predictive capabilities in the preclinical setting [24, 25]. However, the translation of DRP models from preclinical to clinical contexts remains underexplored. In [3], semi-supervised learning with domain generalization losses to generalize DRPs to new data sources. Ma et al. employed pretraining with meta-learning of different contexts and subsequent fine-tuning with few-shot learning to an unseen context to improve the ability of the model to generalize DRPs to new tissues or data sources [12]. In [26, 10, 27], they leveraged DNNs to learn a latent embedding of GEx data, which was utilized either solely in one-model-per-drug response prediction [10, 26], or combined with drug chemical features in a model-for-all-drug manner [27]. Our work aims to address various aspects of DRP with latent gene expression embeddings by enhancing GEx representation learning, emphasizing gene contributions in DRP, incorporating drug chemical features, and training a unified DRP model for a large number of compounds.

In the following sections, we outline the methodology of GExMix, delve into its implementation details, introduce the dual-objective designed to promote the contribution of GEx in DRP, and discuss the use of the learned representation for DRP in the training-data domain along with its transferability to the uncharted domain of human tumors.

## 3 Datasets

### Gene expression data

The gene expression data used in this study includes the following datasets from two sources: 1) the Cancer Cell Line Encyclopedia (CCLE^1^), including 1,247 cell lines covering 35 different cancer types [28]. 2) The Cancer Genome Atlas (TCGA^2^) containing 11, 246 human tumor samples with molecular profiles across 33 cancer types. To optimize the feature set for the model, we selected the top variable 1024 genes, i.e., with the highest standard deviation across all samples.

Our study focuses on the bulk RNA sequencing data, which reflects the summarized gene expression level of tissue sections and the corresponding metadata such as cancer type and subtype, primary tumor site etc. All GEx data have undergone standardization through transcripts per million bases for each gene, followed by a log transformation [29]. Furthermore, GEx data has undergone the Z-score normalization (`scipy.stats.zscore`) to keep the magnitude of all samples on the same scale and mitigate biases and convergence problems introduced by differences in the magnitude of the data.

### Drug screening data

For the DRP task, we leverage the Genomics of Drug Sensitivity in Cancer (GDSC^3^), which has provided an expanded view of pharmacological profiles of *>* 400 compounds across 1, 001 cancer cell lines [8].

### Preprocessing of GDSC drug screen data and TCGA clinical data

For DRP training we only kept drugs tested in more than 10 cell lines in the GDSC dataset. To represent the drug features, we extracted the SMILES (Simplified Molecular Input Line Entry System) [30] notations using PubChempy^4^ for the given drug names. SMILES contain the chemical formula of compounds, and they were tokenized by `tf.keras.preprocessing.text.Tokenizer` into vectors with a maximum of 230 dimensions with zero padding. Notably, the exploration of different text tokenizers remains out of the scope of this study. This is the input to the encoder_*d*_ in Fig. 1.

**Figure 1.**
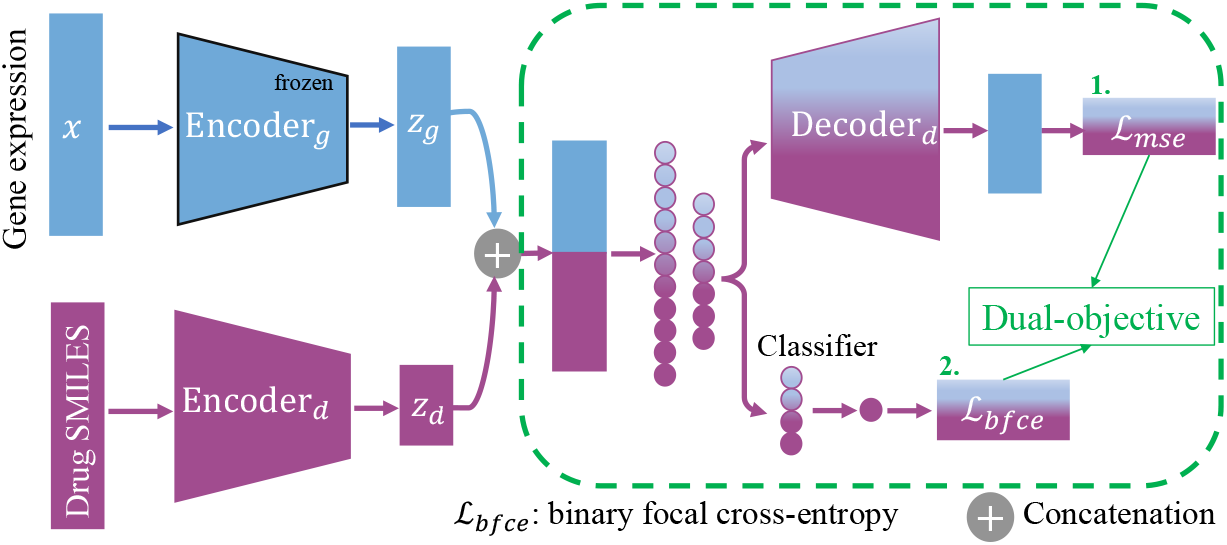
Overview of the proposed framework. A pretrained encoder with GExMix-enriched training set is frozen in dual-objective DRP training. The decoder_*d*_ is used only to reconstruct gene expression latent embeddings.

Drug response in the GDSC dataset is measured as IC_50_, which represents the drug concentration at which relative cell viability is reduced by half [31]. They were binarized [32] to unify the drug response targets across cell lines and human tumors, for which only binary indication is provided. The clinical TCGA data was obtained using the TCGAbiolinks R package [33]. The ‘measure_of_response’ field of patients with drug treatment information was used to assign binary drug responses by labeling complete or partial responses as sensitive and remaining events as resistant. Drug names from both GDSC and TCGA were harmonized with PubChem and ChEMBL notations using the webchem R package [34] for SMILES extraction.

## 4 Methods

In this section, we describe our proposed method, GExMix, a data augmentation for enhancing GEx representation learning in an encoder-decoder model. In subsection 4.2, we show the simple yet effective dual-objective training that promotes focus on gene expression for DRP.

### 4.1 Mixup for Gene Expression

Consider the GEx dataset with samples denoted as *X*_*i*_ ∈ ℛ^*N*^, where *N* is the number of genes in each sample. Each sample is provided with a cancer type *y* in one-hot encoding. The original Mixup [13] suggests combining pairs of samples, such as the *i*-th and *j*-th samples, within their input feature space as well as the label space during training. These pairs of samples are randomly generated and then mixed with a weight that is drawn from the Beta distribution. The mixing process is given by:

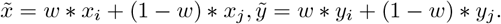

Here, 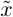 is the augmented samples *x*_*i*_ and *x*_*j*_ with a mixing weight *w*. Likewise, 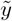 is the resulting label for 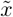.

In GExMix, we extend the basic principles of Mixup to encompass the mixing of samples across multiple data sources. Additionally, we perform per-cancer-type augmentation, where we generate a fixed number of samples for each cancer, to account for the imbalance of the number of different cancer types.

Mixing between different data sources encourages the model to make predictions focusing more on source-invariant features and striping away spurious correlations. This makes the model more robust to covariate shifts between different datasets.

The detailed GExMix is summarized in Algorithm 1. Note that, in the algorithm, we showcase mixing between two samples as an example by setting the shape of W_*mix*_ to have two columns. This illustration is easily extendable to scenarios involving more samples. The mixing weights are drawn from a Beta(*α*) distribution independently for each column. Subsequently, these weights are sorted in row-wise descending order and followed by a row-wise normalization to probabilities (Alg. 1, line 7). The rationale for the descending order is to ensure that the first column gets the highest weights, thereby preserving the cancer types. Lines 12 to 16 implement the stratification across various sources, accounting for the case where one source may be missing. In such cases, samples available in the missing source, e.g., X(*s* = *s*2), regardless of the cancer type, will be used as substitutes. Line 17 shows the linear combination of element-wise multiplication of the weights and samples of X_*mix*1_, and X_*mix*2_.

### 4.2 Gene-Centric Drug Response Prediction

In the DRP task, we freeze the weights of the trained encoder models from the previous section and only utilize them to provide latent embeddings of cell line GEx data.

Figure 1 depicts that the tokenized SMILES annotations are the input to an encoder_*d*_ to reduce the dimension of drug tokens with a classic multi-layer perception (MLP) model with the structure [256 units (Scaled Exponential

#### Algorithm 1

GExMix Data Augmentation

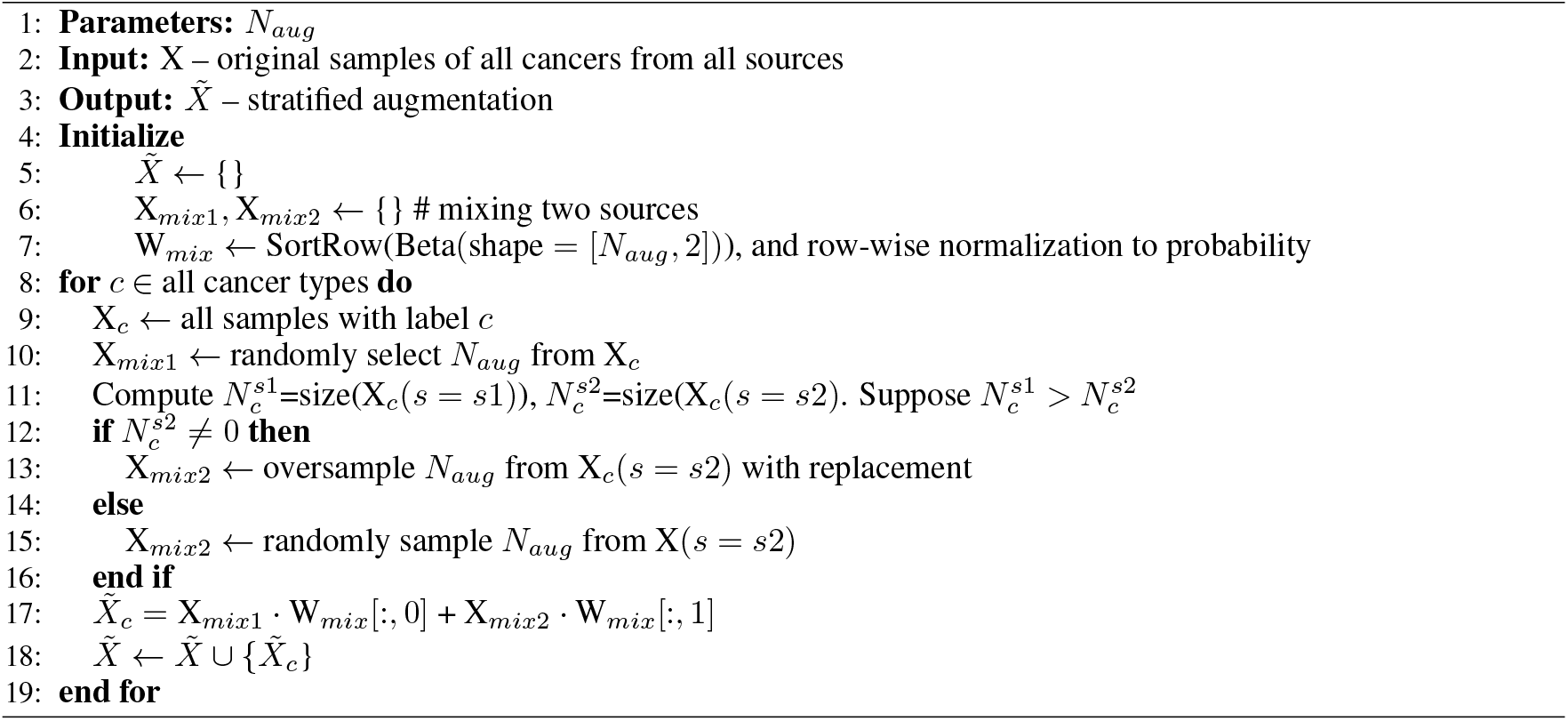

Linear Unit (SELU) activation, BatchNormalization, dropout 0.1) – 6 units – SELU activation – BatchNormalization – dropout 0.1]. Effectively, the tokenized drug features are reduced to 6-dimensional embedding vectors, *z*_*d*_. Then, *z*_*d*_ is concatenated with the extracted latent embeddings of the GEx data, *z*_*g*_. The concatenated embeddings are the input to the dual-objective DRP training, which includes 1) a decoder_*d*_ to reconstruct the concatenated embeddings, and 2) a feed-forward classifier to output the prediction for sensitivity and resistance labels.

The proposed dual-objective consists of the following two losses.

1) Binary focal cross-entropy loss (bfce): To compensate for the underrepresented cell lines that are sensitive to drugs, we adopt the focal cross-entropy loss.

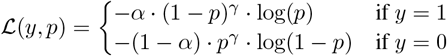

where the value *α* is taken as the frequency of the dominant class in order to give higher weight to the minority class, which is the set of sensitivity drug response labels.

2) Reconstruction loss (mean squared error) of the concatenated gene and drug embeddings: this is the key of the proposed gene-centric training 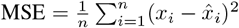.

The dual-objective loss is thus a weighted sum of the above-mentioned losses: ℒ_dual_ = *w*_bfce_ ·ℒ_bfce_ + *w*_MSE_ · ℒ_MSE_. We take a uniform distribution between *w*_bfce_ and *w*_MSE_, thus 0.5, as suggested in [35, 36].

### 4.3 Training Procedure

#### Gene expression encoder

To apply GExMix for gene expression data representation learning, we adopted the encoder from [10], CODE-AE. We considered two variants: CODE-AE-Base and CODE-AE-Adv, where “Adv” refers to the adversarial loss used to align TCGA and CCLE. For an in-depth understanding of detailed encoder design, we direct the readers to [10]. The key proposition of CODE-AE models is to learn a shared encoder for GEx data from two data sources, i.e., TCGA and CCLE, such that source-dependent features are maximally removed from the shared encoder and kept in the source-specific encoder. Simultaneously, the model aims to maximize the overlap of latent distributions between two sources. As mentioned, CODE-AE-Adv enforces indistinguishability between the concatenation of the source-specific and shared embeddings from either source.

Additionally, we implemented a *β*-VAE model [37] for GEx representation learning as a baseline model. Here *β* is introduced to control the degree of disentanglement of the latent representation while trading off the reconstruction quality. For the VAE model, we adopted a two-layer MLP design for the encoder and a symmetric decoder, i.e., the encoder consists of 256 and 64 units in each layer, encoding the input into 36 latent dimensions. The decoder then reconstructs the 36 latent dimension embeddings through 64 and 256 units back to the same shape as the input. We chose a *β* value of 0.01 for the KL divergence. It has been shown that with the same bottleneck dimension, a higher *β* yields worse reconstruction [37]. The choice of 0.01 is to improve the reconstruction w.r.t. the standard VAE, while still preserving a relatively continuous distribution of the latent space.

Models were trained with GExMix enhanced training set at a starting learning rate of 0.01, and a `ReduceLROnPlateau` callback in `Tensorflow` with `factor=0.5, patience=3, min_lr=0.0001`. In other studies, the training set consists only of augmented samples [13, 38]. However, in our experiments, we included the original data since a deteriorated performance was observed otherwise. The training of the VAE model continued until there was no improvement in the total loss on the validation set for five consecutive epochs.

The metrics used to evaluate performance are the *R*^2^ coefficient for the reconstruction quality. *R*^2^ is a measure of how much variability of the data can be explained by the model, computed by 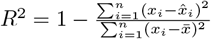. Here *x*_*i*_ is the *i*-th feature in the GEx input, and 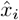 is the reconstructed feature.

#### Drug response classifier

We performed five-fold cross-validation on the CCLE-GDSC data. In each fold, the training and validation data were derived from an 80-20% random split of all existing cell line-drug pairs from CCLE-GDSC data, with the additional TCGA drugs response as withheld test set. Note that the trained DRP model with CCLE-GDSC is directly tested on the TCGA dataset without further fine-tuning. This is confronted by inherent differences 1) in GEx profiles of samples due to biological and technical confounders [10], 2) between drug response labels of cell lines and patient tumors, and 3) the binarization thresholds for drug response are substantially misaligned between preclinical and clinical applications.

The DRP model was trained for 10 epochs and tested on the validation set. The parameters for the binary focal loss, i.e., *γ* and *α* were set to be 10 and 0.35, respectively, suggested by the hyperparameter exploration with the Optuna library^5^. The models were implemented with tensorflow keras version 2.10.0.

The evaluation metric for DRP was the area under the Receiver Operating Characteristics curve (ROC-AUC) for each drug across all drugs on the test set. Specifically, for TCGA drugs, we only considered drugs with more than 10 unique patients, resulting in 24 valid drugs.

## 5 Results

We perform ablation studies for two key components of our proposed framework: GExMix, a data augmentation method, and a gene-centric dual-objective DRP training. The overall performance is summarized in Table 1. Specifically, we pretrain three encoder-decoder models with the GEx data from TCGA and CCLE, i.e., a *β*-VAE, CODE-AE-Base, and CODE-AE-Adv [10]. Subsequently, the trained encoders are utilized for DRP as described in Fig. 1. The trained DRP models are evaluated on the CCLE and the TCGA validation sets, and organized into “Single-objective for DRP” and “Dual-objective for DRP” columns in Table 1.

**Table 1:** Model performance with ablation studies. Here, the performance is reported on five-fold cross-validation sets. DRP: drug response prediction. DA: GExMix data augmentation with 1000 samples per cancer type. Note that we compute the ROC-AUC for each drug, then average across all drugs. Values are shown in mean *±* std.

| Models | Reconstruction $R^2$ | Single objective for DRP | | Dual objective for DRP | |
| --- | --- | --- | --- | --- | --- |
|  | TCGA+CCLE val set | CCLE | TCGA | CCLE | TCGA |
| VAE | $0.789 \pm 0.002$ | $0.665 \pm 0.106$ | $0.473 \pm 0.087$ | $0.563 \pm 0.084$ | $0.510 \pm 0.069$ |
| CODE-AE-Base | $0.911 \pm 0.008$ | $0.736 \pm 0.154$ | $0.461 \pm 0.116$ | **$0.731 \pm 0.151$** | $0.460 \pm 0.110$ |
| CODE-AE-Adv | $0.876 \pm 0.005$ | $0.726 \pm 0.157$ | **$0.531 \pm 0.112$** | $0.725 \pm 0.155$ | $0.521 \pm 0.117$ |
| VAE+DA | $0.792 \pm 0.004$ | $0.690 \pm 0.123$ | $0.509 \pm 0.066$ | $0.569 \pm 0.067$ | $0.512 \pm 0.065$ |
| CODE-AE-Base+DA | $0.988 \pm 0.008$ | $0.729 \pm 0.151$ | $0.508 \pm 0.099$ | $0.716 \pm 0.151$ | $0.554 \pm 0.090$ |
| CODE-AE-Adv+DA | $0.978 \pm 0.007$ | **$0.737 \pm 0.152$** | $0.517 \pm 0.100$ | $0.724 \pm 0.149$ | **$0.561 \pm 0.104$** |

### 5.1 GExMix Data Augmentation Ablation Experiments

Focusing on the reconstruction quality of models with “+DA” (data augmentation) and without, we observe consistent improvements of R^2^ coefficients on the validation sets across all models. This is particularly apparent in two CODE-AE models with the R^2^ improved from 0.911 to 0.988 for CODE-AE-Base and from 0.876 to 0.978 for CODE-AE-Adv.

Comparing models in the single-objective DRP tests, with and without data augmentation, we observe the VAE demonstrates improvement in both CCLE and TCGA DRP performance, however, a competition between the CCLE and TCGA performance is observed for the two CODE-AE variants: CODE-AE-Adv+DA shows increased performance in CCLE drugs but a decrease in TCGA drugs, while CODE-AE-Base+DA exhibits the opposite trend. This could suggest that high performance on CCLE might indicate a risk of overfitting the training data distribution, which may lead to reduced transferability. The observed discrepancy in CCLE and TCGA test sets between VAE and CODE-AE models might suggest that a robust representation is pivotal for achieving high DRP performances. Thus, further research is needed to fine-tune parameters for the VAE to achieve comparable performance with the CODE-AE model while balancing the *β* and the latent dimensions.

### 5.2 GExMix Effect on Latent Representation

To visually demonstrate the effect of the proposed GExMix method on GEx representation, we apply UMAP visualization to the learned latent representation with and without GExMix, shown in Fig. 2C and B, respectively. As a comparison, the UMAP of the original data is shown in Fig. 2A (1024 gene features). Given that the original CODE-AE models have been designed explicitly to align the distribution of CCLE and TCGA in the latent space, to specifically assess the impact of the proposed GExMix, we implement a *β*-VAE to learn the latent GEx representation.

**Figure 2.**
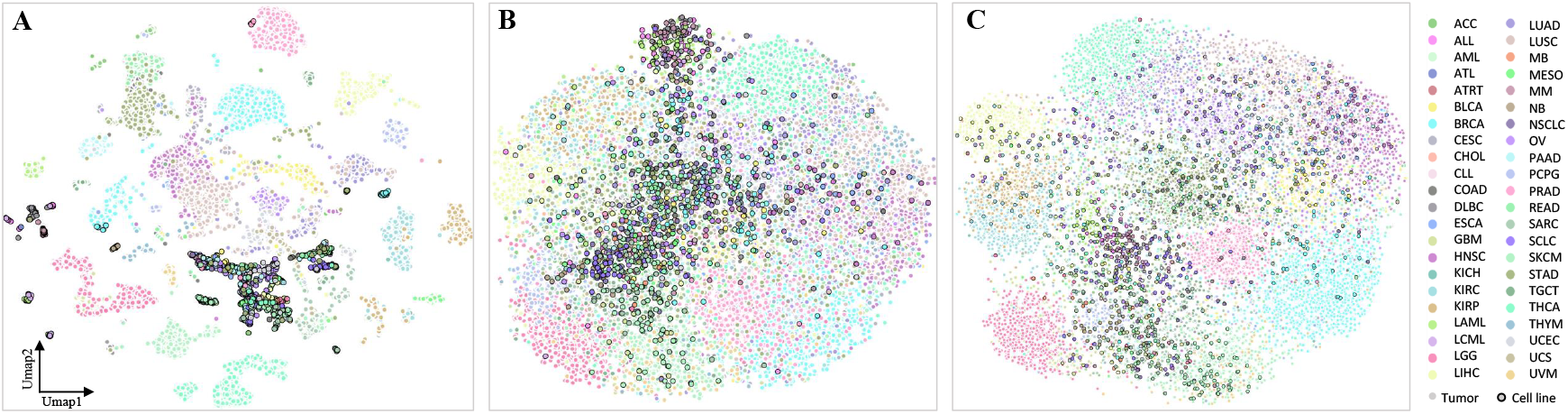
UMAP visualization of the tumor and cell line samples. A: The original features. B: The learned latent representation without data augmentation. C: The learned latent representation with data augmentation. The samples include 10,514 tumor samples (white edge color dots) and 1,107 cell lines (black edge color dots), covering 44 cancer types. The models used in B and C are standard *β*-VAEs, where *β*=0.01 for encoding the GEx data.

Shown in Fig. 2A, the GEx profile of cell lines is inherently different from that of human tumors, resulting in isolated clusters. The use of *β*-VAE learning with the original data increases the alignment of cell lines to their corresponding tumor representations, shown in Fig. 2B. This effect is further amplified by adding 1000 augmented samples per cancer type to the original training set, as shown in Fig. 2C.

### 5.3 GExMix Hyperparameter Effect

Here, we show how the reconstruction quality on the validation set is affected by different augmentation methods, i.e., GExMix and random, as well as the number of augmented samples per cancer type.

In Figure 3, we show that our proposed stratified augmentation leads to an improvement in R^2^ values on the validation set. This becomes more pronounced as the number of augmented samples increases and peaks around #augment = 2000.

**Figure 3.**
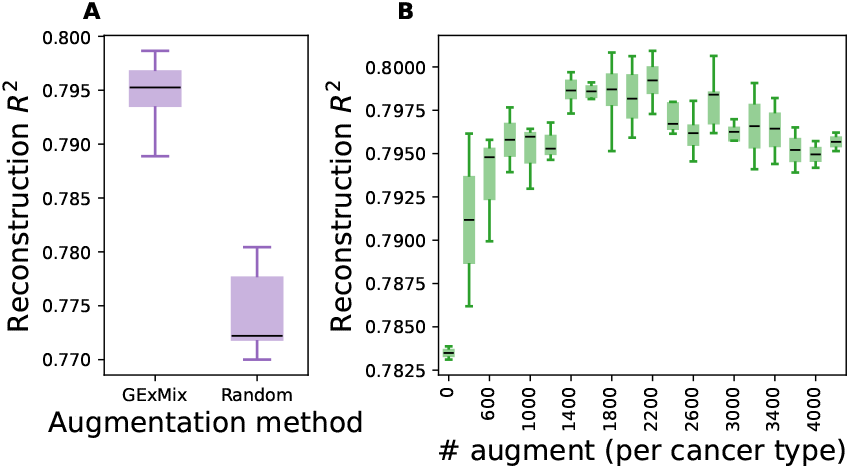
How the reconstruction quality, measured by *R*_2_ coefficient is affected by (A) stratified augmentation as in GExMix, or random augmentation, and (B) the number of augmented samples. A and B are obtained through hyperparameter tuning in 50, and 200 trials, respectively.

The reconstruction quality plateaus or even deteriorates with a further increase in augmented samples. The results are derived from 50 random trials within the given set of parameter options, i.e., #augment ∈ [0, 4500] with a step-size of 300, and the augmentation method ∈ [GExMix, Random] with #augment=500. The error bars denote the standard deviation across trials with the same configuration.

### 5.4 Dual Objective Ablation Experiments for DRP

Performances of ablation experiments w.r.t. dual-objective training are organized into “Single-objective for DRP” and “Dual-objective for DRP” columns in Table 1. We take the encoders from previously pretrained models for DRP without fine-tuning the encoders’ weights. The purpose is to assess the quality of the learned GEx representations for the DRP task.

#### The dual-objective training prevents overfitting to CCLE drugs but heavily relies on the quality of learned GEx representation

When comparing models **without** GExMix in both single- and dual-objective columns, we note that the dual-objective training does not yield improved performance for two variants of CODE-AE models. It only exhibits benefits in translating to TCGA for the VAE model, albeit at the cost of degraded performance on CCLE drugs. This underscores the importance of initially learning a high-quality GEx representation.

#### The dual-objective training improves the DRP transfer to TCGA drugs, leveraging the enhanced GEx representation with data augmentation

In the comparison of model **with** GExMix in single- and dual-objective training columns, it is observed that adding the dual-objective often leads to a slight performance decrease on CCLE drugs, consistent with the previous observation. However, the dual-objective demonstrates significant improvement in the TCGA DRP performance for two CODE-AE models, with *p*-values *<* 0.0007. Notably, the model CODE-AE-Adv+DA+Dual achieves the best performance on the TCGA drugs. This underscores the benefit of the GExMix in enhancing the GEx representation, which subsequently benefits the gene-centric DRP.

The per-drug ROC-AUC distributions for both CCLE and TCGA drugs are depicted side-by-side for each model in Fig. 4. These models consist of three baseline models, VAE, CODE-AE-Base, and CODE-AE-Adv, and their counterparts with the proposed data augmentation and the dual objective “+DA+Dual”. Statistical tests (one-sided Wilcoxon signed-rank tests) of the ROC-AUC distributions for TCGA drugs reveal significant improvement in the two model pairs, i.e., (VAE, VAE+DA+Dual), and (CODE-AE-Base, CODE-AE-Base+DA+Dual). The *p*-values for these two pairs are 0.028 and 0.008, respectively. The *p*-value for the pair (CODE-AE-Adv, CODE-AE-Adv+DA+Dual) is 0.08 (not significant), which could result from the explicit integration of source aligning in the encoder of CODE-AE-Adv itself between cell lines and human tumors. As demonstrated in Fig. 2, GExMix promotes the alignment between two sources, leading to a significant improvement for VAE and CODE-AE-Base. However, the performance gain for CODE-AE-Adv is less pronounced. This confirms the source alignment between cell lines and tumors is crucial for knowledge transferring for DRP, as proposed in [10]. Overall, CODE-AE-Adv+DA+Dual achieved the best performance in TCGA drugs, boosting the number of TCGA drugs with a ROC-AUC *>*0.6 from 8 (in CODE-AE-Adv) in the single objective DRP to 11 out of 24 drugs.

**Figure 4.**
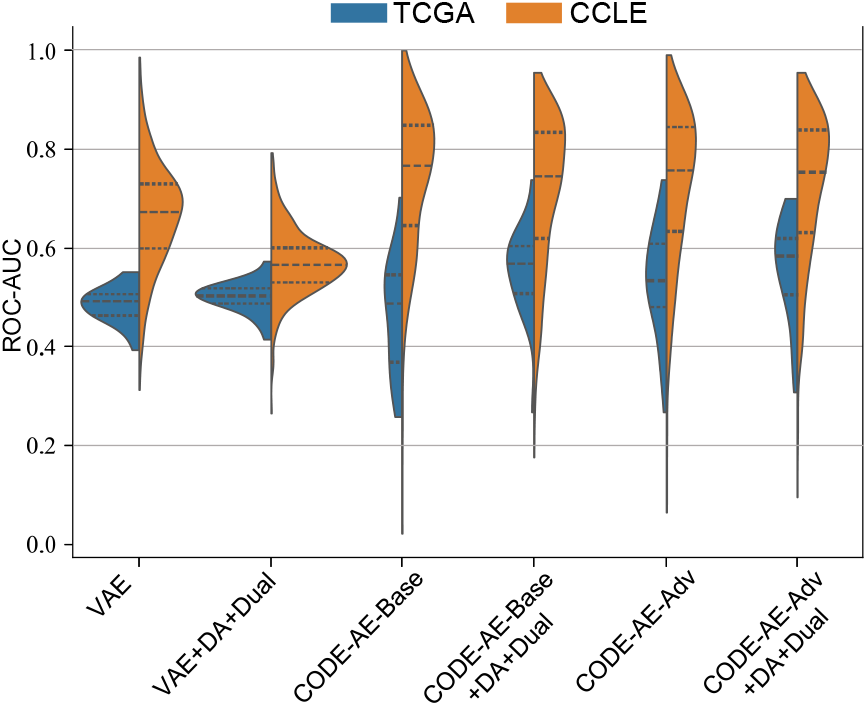
Model performances measured as ROC-AUC per drug for all drugs within TCGA (blue) and CCLE (orange) datasets. Only drugs with *>* 10 samples that have both sensitive and resistant samples were considered. DA: data augmentation

Figure 5 further illustrates enhanced predictions of drug response in a few selected drugs with our proposed method. These drugs are determined by taking the union of top-performing drugs across models for CCLE drugs (Fig. 5A) and TCGA drugs (Fig. 5B), respectively. While models with GExMix and the dual-objective training exhibited comparable performances for CCLE drugs, our top-performing model (CODE-AE-Adv+DA+Dual) showed elevated performances for TCGA drugs. Specifically, the performance enhancement is particularly pronounced for chemotherapeutic compounds such as methotrexate, 5-fluorouracil, cisplatin, and epirubicin in the TCGA dataset. This highlights the DRP generalization abilities of our proposed integration of data augmentation and gene-centric training strategies.

**Figure 5.**
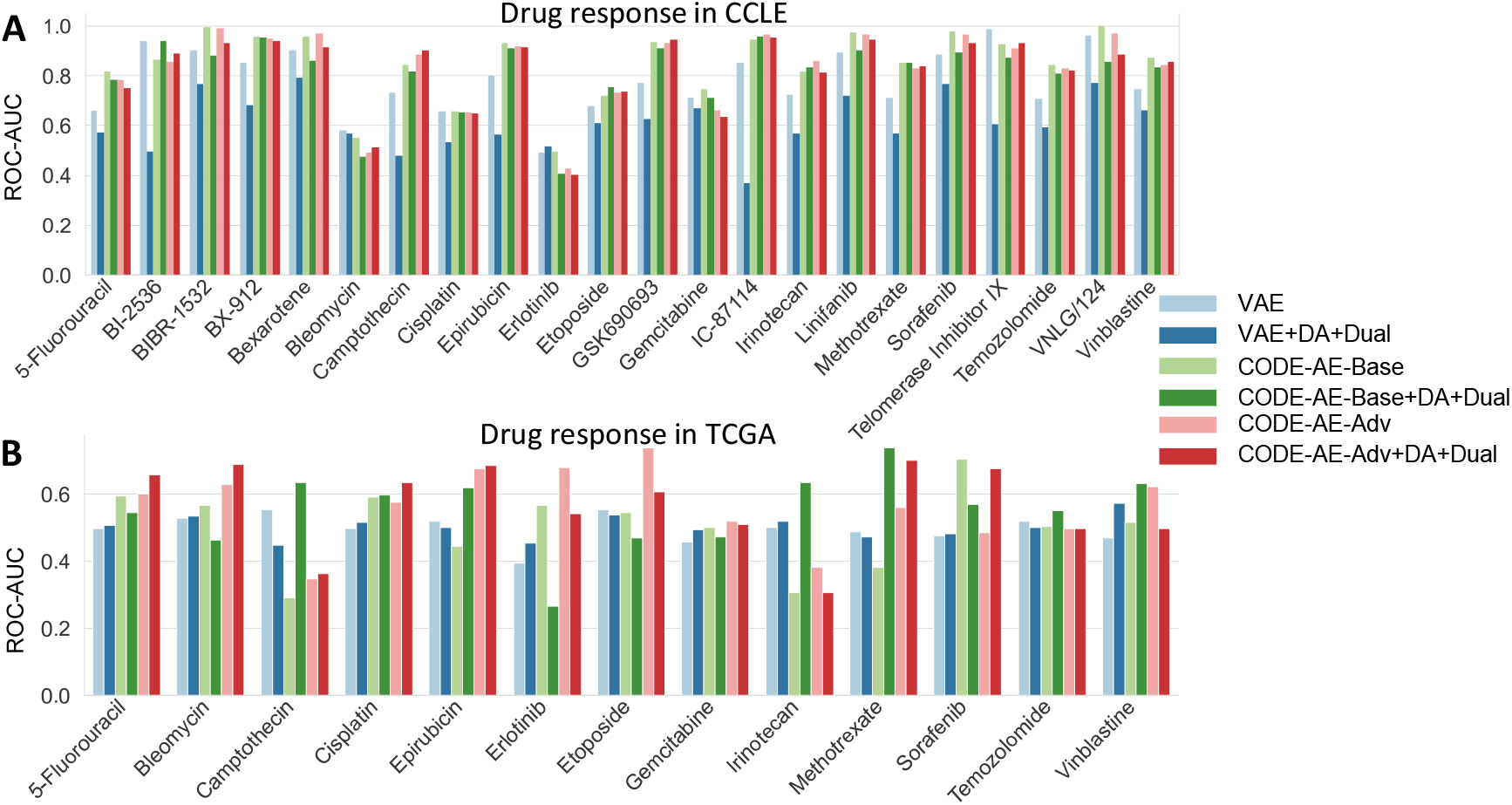
The union of top performing drugs across models in CCLE (A) and TCGA drugs (drugs with *>* 10 unique samples) (B). DA: data augmentation. Dual: with dual objective training for DRP.

## 6 Conclusion

In conclusion, our proposed data augmentation, GExMix, presents a promising tool to enhance gene expression representation learning, addressing challenges associated with GEx data representation for precision medicine. GExMix strategically employs data augmentation to compensate for imbalances in cancer types and data sources, thereby enabling seamless integration across diverse datasets and providing promising avenues for improving representation learning with GEx data.

The introduction of a dual-objective training scheme, accentuating the contribution of the latent representation of GEx data, demonstrates great potential in leveraging molecular features together with drug chemical features for DRP, paving the way for precision medicine.

Our experiments validate the benefit of GExMix in improving the latent representation of GEx data in all tested models. Moreover, the integration of GEx data from diverse sources, coupled with gene-centric dual-objective training, leads to an improved transferability from cell lines to human tumors, showcasing the efficacy of our approach with untapped potential for extension to other omics beyond GEx data in cancer research.

### Limitations and future work

Despite the contribution, we have identified a few limitations in our work, which we believe hold opportunities for future research and development.

Our experiments were conducted with only two publicly available datasets on cancer and trained with an unsupervised fashion, i.e., not incorporating cancer labels into the representation learning. blackOur preliminary results showed that the supervised objective yields no benefit on DRP. In the future, we plan to explore multiple dataset integration and investigate the effect of cancer-type supervision on DRP. Potentially, other omics data beyond GEx could be fused with GExMix. Furthermore, GEx data from any disease or condition can be added to enhance the overall GEx representation.

Regarding the DRP, further exploration of alternative characterization of drug chemical features other than SMILES could be explored and could potentially provide additional insights.

Moreover, to reveal biological insights and predictive biomarkers from these models, future work should focus on the interpretation of 1) the latent dimensions with biological information, such as gene pathways, disease biomarkers, etc., and 2) gene-drug interactions w.r.t. the drug response. Furthermore, applying GExMix to explicitly enrich DRP variance coverage presents an intriguing avenue for future investigations.

## Footnotes

1 https://depmap.org/portal/download/all/

2 https://www.cancer.gov/ccg/research/genome-sequencing/tcga

3 https://www.cancerrxgene.org/downloads

4 https://pubchempy.readthedocs.io/en/latest/

5 https://optuna.org/

